# Ageing disrupts reinforcement learning whilst learning to help others is preserved

**DOI:** 10.1101/2020.12.02.407718

**Authors:** Jo Cutler, Marco Wittmann, Ayat Abdurahman, Luca Hargitai, Daniel Drew, Masud Husain, Patricia Lockwood

**Affiliations:** Department of Experimental Psychology, University of Oxford, United Kingdom; Wellcome Centre for Integrative Neuroimaging, Department of Experimental Psychology, University of Oxford, United Kingdom; Centre for Human Brain Health, School of Psychology University of Birmingham, United Kingdom; Department of Psychology, University of Cambridge, United Kingdom

**Keywords:** Prosocial behaviour, ageing, reinforcement learning, computational modelling

## Abstract

Reinforcement learning is a fundamental mechanism displayed by many species. However, adaptive behaviour depends not only on learning about actions and outcomes that affect ourselves, but also those that affect others. Here, using computational reinforcement learning models, we tested whether young (age 18-36) and older (age 60-80, total n=152) adults can learn to gain rewards for themselves, another person (prosocial), or neither individual (control). Detailed model comparison showed that a model with separate learning rates for each recipient best explained behaviour. Young adults were faster to learn when their actions benefitted themselves, compared to helping others. Strikingly, compared to younger adults, older adults showed preserved prosocial learning rates but reduced self-relevant learning rates. Moreover, psychopathic traits were lower in older adults and negatively correlated with prosocial learning. These findings suggest learning how to benefit others is preserved across the lifespan with implications for reinforcement learning and theories of healthy ageing.

---

Learning associations between actions and their outcomes is fundamental for adaptive behaviour. To date, the majority of studies examining reinforcement learning have tested how we learn associations between actions and outcomes that affect ourselves, and largely focused on these processes in young age, both in humans and other species^1–5^. However, such self-relevant learning may be computationally separable from learning about actions that help other people. Studies suggest slower learning of associations between actions and outcomes when they are about^6^ or affect others^7^, henceforth referred to as ‘prosocial learning’.

Senescence is associated with a multitude of changes including declines in cognitive functioning and perception, but perhaps preservation of affective processing and social cognitive abilities^8–10^. However, less is known about how ageing affects social behaviour, despite the critical importance of this question. Social isolation has been found to be as damaging to physical health as smoking or excessive drinking^11^. Social behaviours that benefit others – prosocial behaviours – are vital for maintaining social bonds and relationships^12^ across the lifespan. In addition to the benefits for others, prosociality has been linked to improved life satisfaction^13^, mental wellbeing^14^, and physical health^15^ for the person being prosocial, all of which could contribute to healthy ageing. A key aspect of prosocial behaviour is the ability to learn associations between our own actions and outcomes for other people^7^. Here, we use computational models of reinforcement learning in young and older participants to examine the mechanisms that underpin self-relevant and prosocial learning and associations with healthy individual differences in socio-cognitive ability.

Reinforcement Learning Theory (RLT) provides a powerful framework for understanding and precisely modelling learning^16^. In RLT, prediction errors signal the unexpectedness of outcomes and affect the choices we make in the future. The influence that prediction errors have on choices can be modelled individually through the learning rate, which quantifies the effect of past outcomes on subsequent behaviour. The plausibility of reinforcement learning as a core biological mechanism for action-outcome associations is bolstered by our understanding of neurobiology, with prediction errors encoded by single neurons in the ventral tegmental area^17^.

Although essential for successful adaptive behaviour, several studies suggest that our propensity for reinforcement learning declines in later life^10^. Compared to younger adults, older adults show learning impairments particularly when action-outcome associations are probabilistic^18^ or reverse^19^. Age-related declines in learning ability have been linked to functional and structural changes in frontostriatal circuits^20,21^ and dopamine transmission^18,22^, which shows a significant age-related decrease^23–25^ and has a key role in coding prediction errors^2,26,27^. Indeed, one study showed that administering L-DOPA, a dopamine precursor, to older adults increased their learning rate^28^. Therefore, if reinforcement learning in general declines in older age, we would hypothesise lower learning rates for both self-relevant and prosocial learning in older, compared to younger, adults.

Alternatively, prosocial learning may depend not only on our learning ability, but also our motivation to help others. Results from experiments using economic games to measure prosociality have found that older adults tend to be more generous^29,30^. There is also evidence of an age-related increase in charitable donations to individuals in need^31^. At work, older adults engage in more prosocial behaviours than younger adults, according to both self-report data and colleagues’ ratings^32^. Finally, self-reported altruism and decisions to donate to others have been shown to increase with age^33^. However, one limitation of these studies is that the paradigms often place self and other reward preferences in conflict. Money for the other person depends on less money for oneself. Moreover, older adults generally have higher accumulated wealth, which would be an important confound in studies of monetary exchange^34^. Prosocial learning avoids this confound by separating outcomes for oneself from outcomes for others. If older adults do indeed value outcomes for others more than young adults, we might expect that whilst self-relevant learning declines with ageing, prosocial learning could be preserved. Comparing young and older adults on self-relevant and prosocial learning provides an opportunity to dissociate how possible age-related changes in cognitive ability and social behaviour impact on learning.

While studies point to potential group differences between young and older adults, there is also substantial variability in self and other reward sensitivity. For example, psychopathy is a key trait associated with decreased prosocial behaviour and altered self and other reward processing^35,36^. Psychopathy has dysfunctional affective-interpersonal features at its core^37,38^ but is also characterised by lifestyle and antisocial traits^39^. At the extreme, psychopathy is a severe personality condition linked to poor life outcomes, violence, and criminality^40–42^. However, several studies suggest similar behavioural and neural profiles between community samples with high levels of psychopathic traits^43^ and those with clinical diagnoses of psychopathy, consistent with the Research Domains of Criteria (RDoC) conceptualisation of a dimensional approach to psychiatry^44^. This RDoC approach suggests that psychiatric disorders can be thought of as dimensional, rather than categorical, constructs. In a similar vein, psychopathic traits can be captured on a continuum spanning clinical samples and the general population, with a range of scores on that continuum for healthy people. Self-report measures of psychopathic traits that mirror the latent structure of clinical psychopathy measures, comprising antisocial and interpersonal dimensions, are available to use in samples from the general population^39^.

Intriguingly, preliminary evidence suggests that ageing may also be associated with changes in psychopathic traits, which could have important implications for our understanding of an ageing population. Epidemiological studies show that criminal activity increases during adolescence then declines in older adulthood^45^. Antisocial and aggressive behaviours also significantly decrease in older age, with young adults (age 16-24 years) having the highest rates of homicide^46^. Even within violent male offenders, psychopathic traits linked to an antisocial lifestyle are negatively correlated with age^47^. In community samples, ageing is associated with a decrease in both the antisocial-lifestyle (antisocial and impulsive behaviours) and affective-interpersonal (lack of empathy and guilt) elements of psychopathic traits^48^. These studies highlight the importance of assessing how differences in psychopathic traits could map on to differences in prosocial behaviours. However, no existing work has examined this question.

Taken together, previous research supports opposing hypotheses for how ageing is associated with self-relevant and prosocial reinforcement learning. On the one hand, evidence suggests that older adults should be impaired at learning, regardless of the recipient, consistent with ageing-related declines in learning ability and dopamine transmission. On the other hand, potential increases in valuing outcomes for others in older, compared to younger, adults would predict preserved prosocial learning ability but reduced self-relevant learning ability. Finally, we expected variation in psychopathic traits to be associated with learning for others but not self in both age groups.

To distinguish between these competing hypotheses, we tested 75 young (aged 18-36, mean=23.07, 44 females) and 77 older (aged 60-80, mean=69.84, 40 females) adults carefully matched on gender, years of education, and IQ. Participants completed a probabilistic reinforcement-learning task (Figure 1) designed to separate self-relevant (rewards for self) from prosocial learning (rewards for another person), as well as controlling for the general valence of receiving positive outcomes (rewards for neither self or other).

**Figure 1.**
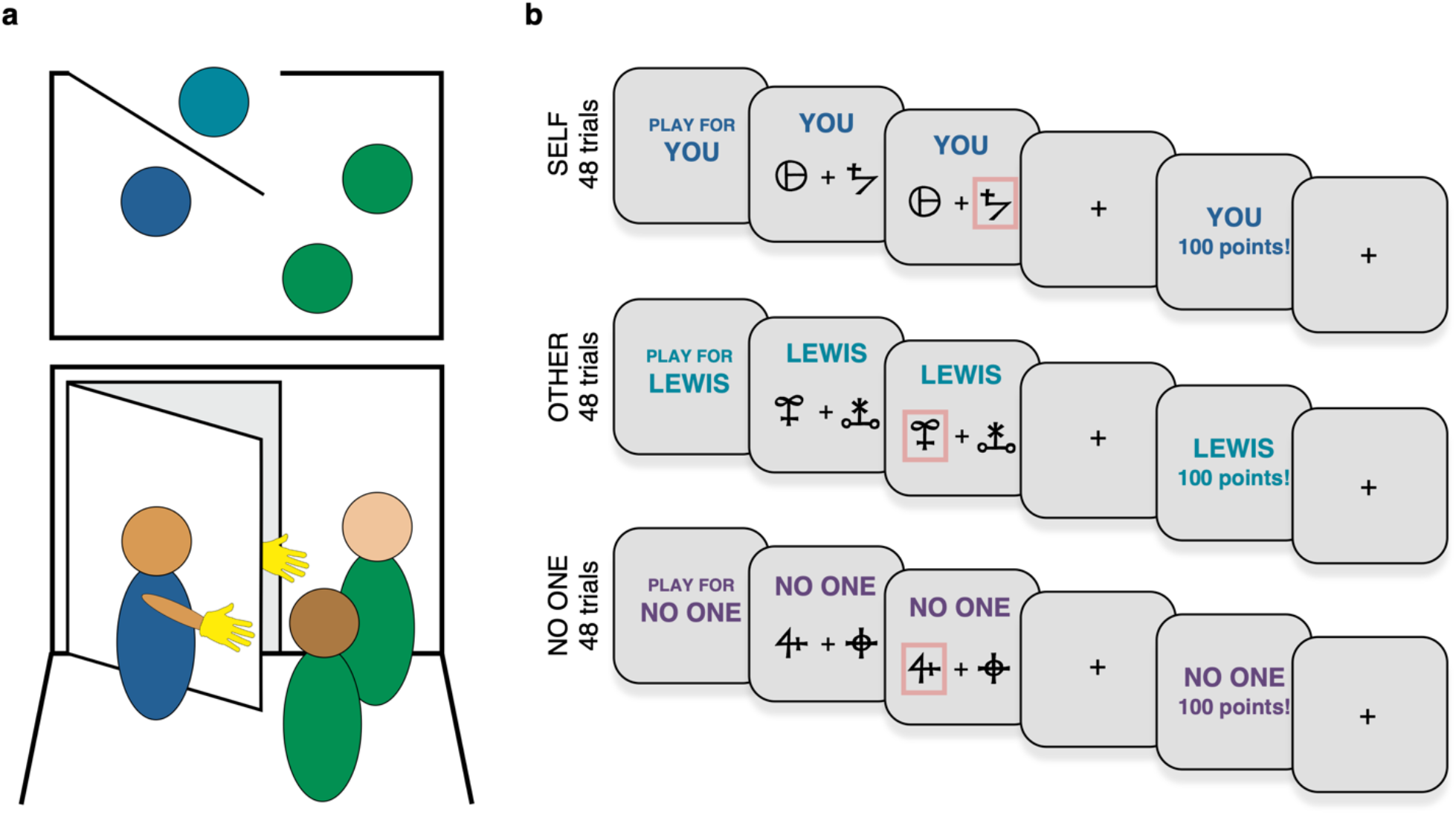
Prosocial learning task and social role assignment. **(a)** The role assignment procedure involved the participant (dark blue), confederate (light blue) and two experimenters (green). Top: from above showing the positioning of the participant and two experimenters inside the testing room, and the confederate the other side of the door. Bottom: the participant and confederate wore a glove to disguise their identity and waved to each other from either side of the door. Participants were instructed that they would be assigned to roles of ‘Player 1’ and ‘Player 2’ but the participant was always assigned to be Player 1. After this procedure participants were informed that they would play a game where they could gain rewards for themselves, the other participant (Player 2) or neither participant. They were told that Player 2 would not play the same game for them and that Player 2 would not know that they may receive an additional bonus based on the choices the participants. This meant that participants’ choices were made anonymously and should not be affected by reputational concerns. **(b)** Participants performed a reinforcement learning task (‘prosocial learning task’) in which they had to learn the probability that abstract symbols were rewarded to gain points. At the beginning of each block, participants were told who they were playing for, either themselves, for the other participant, or in a condition where no one received the outcome. Points from the ‘self’ condition were converted into additional payment for the participant themselves, points from the ‘other’ condition were converted into money for Player 2 and points from the ‘no one’ condition were displayed but not converted into any money for anyone.

Detailed model comparison showed that a computational model with separate learning rates best explained how people learn associations for different recipients (Figure 2). Young adults were faster to learn when their actions benefitted themselves, compared to when they helped others. Strikingly however, older adults showed a reduced self-bias, with a relative increase in the rate at which they learnt about actions that helped others, compared to themselves (Figure 3a & b). Older adults had significantly reduced levels of psychopathic traits compared to younger adults and in older adults, lower psychopathic trait scores correlated with prosocial learning rates (Figure 4a & b). These effects were not explained by individual differences in IQ, memory or attention abilities. Overall, we show that older adults are less self-biased in reinforcement learning than young adults, and ageing is associated with a decline in psychopathic traits. These findings suggest learning how our actions help others is preserved across the lifespan.

**Figure 2.**
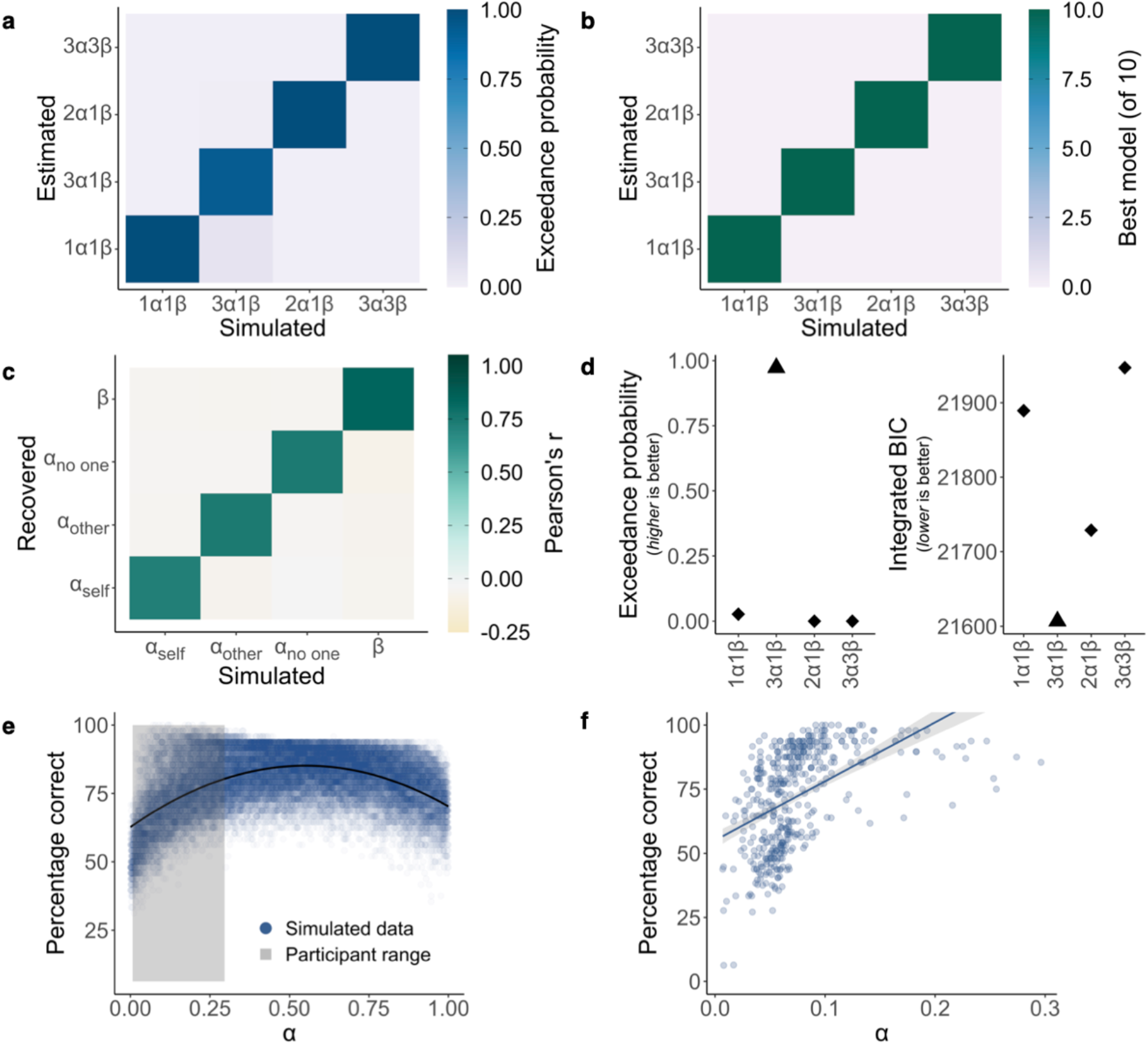
Model identifiability, parameter recovery and simulation. **(a)** Model identifiability average exceedance probability confusion matrix and **(b)** model identifiability best model selection confusion matrix. Data were simulated from 150 synthetic participants with each of our four models then Bayesian model selection was applied, and this procedure was repeated 10 times. Identifiability is shown by strong diagonals. **(c)** Parameter recovery was performed on data simulated by the winning 3α1β model from 1296 synthetic participants. Confusion matrix represents correlations between simulated and fitted parameters. Strong correlations on the diagonal show parameters can be recovered. **(d)** The 3α1β model (▴) is the best model on both exceedance probability and integrated Bayesian Information Criterion (BIC) fit measures. **(e)** Average percentages of correct choices (high probability of reward option) associated with 30,000 simulated α values (10,000 synthetic participants, 3 recipient conditions) show that an optimal learning rate is approximately 0.55 in this task. The range of α values for our participants was below this peak (grey shading), such that a higher learning rate was associated with better performance. **(f)** Correlation between percentage correct and learning rate across participants. There was a significant correlation between learning rate and accuracy (*r*_s(150)_=0.58 [0.46, 0.68], *p*<0.001) (see Supplementary Table 3 for each separate age group and recipient combinati α on; *r*_s_ in all cases > 0.46, *p*s <0.001). Shaded area represents 95% confidence interval.

**Figure 3.**
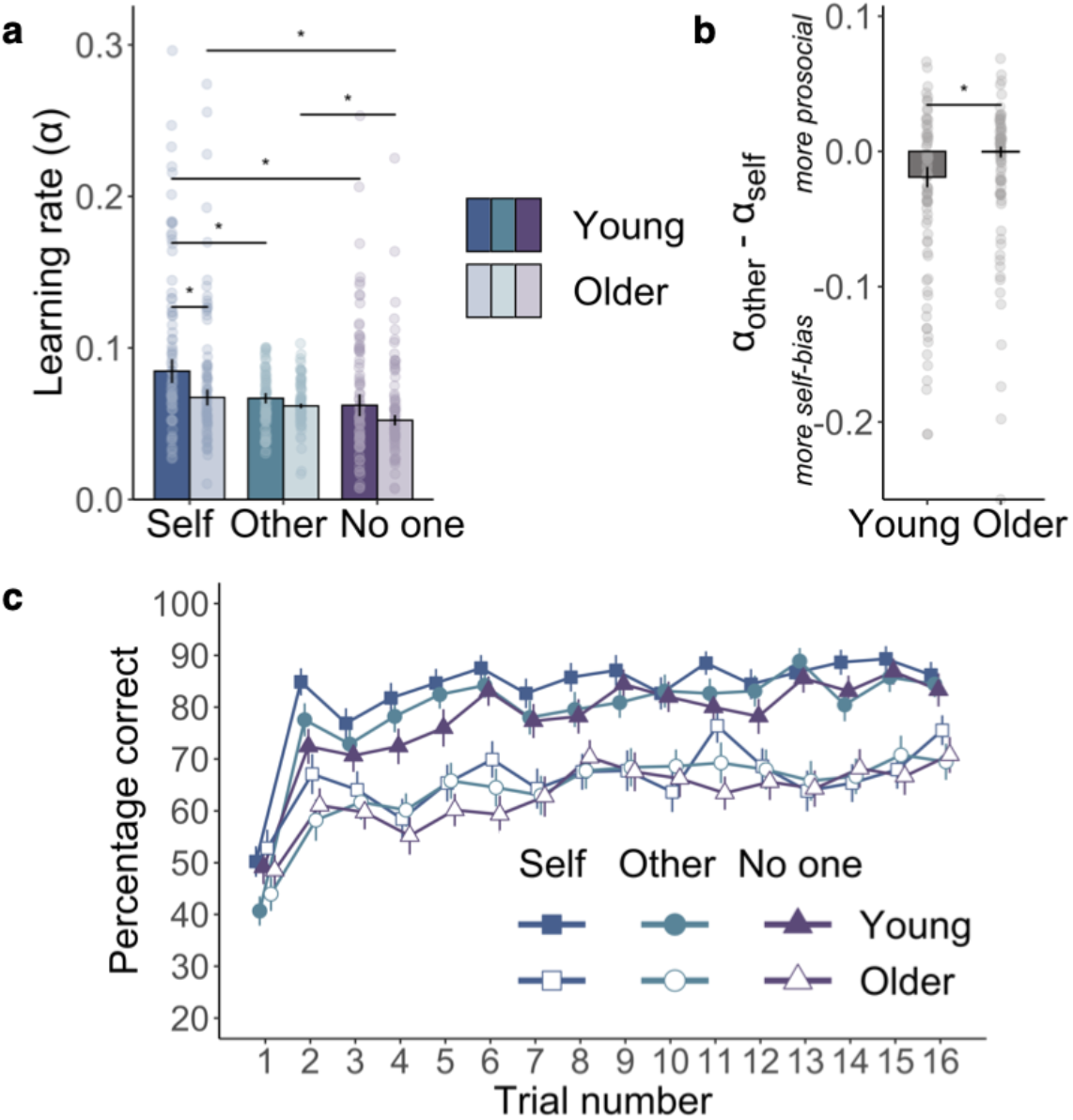
Age-group differences in accuracy and learning rates. **(a)** Comparison of learning rates from the computational model show older adult’s prosocial learning was preserved: learning rates in the other condition did not differ from the self condition or from young adult’s prosocial learning rates. Only young participants showed self-bias, n=150 (75 young, 75 older). Bars show group median, error bars are standard error of the median, asterisks represent significant between-group and within-group Wilcoxon t-tests (*p*<0.05). **(b)** Median difference between learning rates in the other and self conditions illustrates the larger self-bias in young, compared to older, adults, n=150 (75 young, 75 older). Error bars are standard error of the median, the asterisk represents the significant age group * recipient [self vs. other] interaction from the robust linear mixed-effects model (*p*<0.001). **(c)** Group-level learning curves showing choice behaviour in the three recipient conditions for each age group. Trials are averaged over the three blocks (48 trials total per recipient presented in three blocks of 16 trials) for the self, other, and no one recipients, n=152 (75 young, 77 older). Points show group mean, error bars are standard error of the mean.

**Figure 4.**
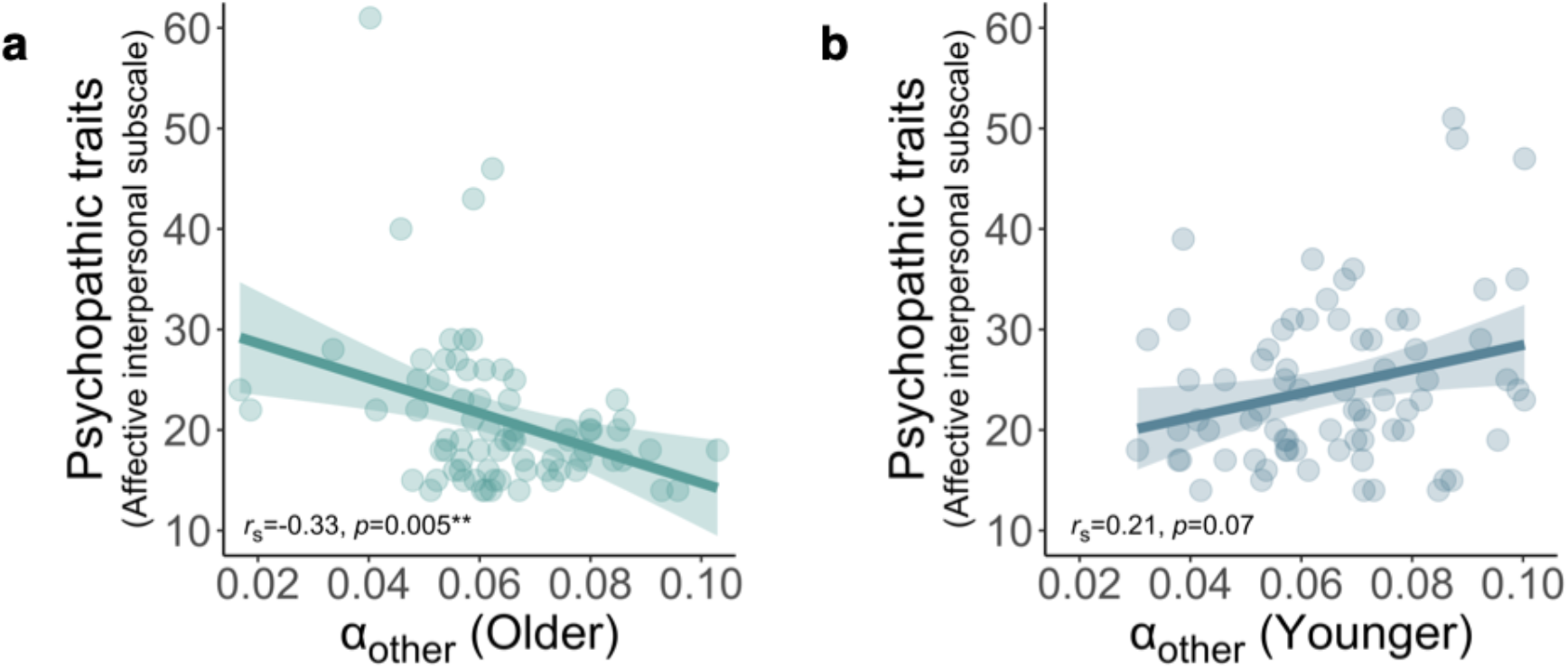
Correlations between prosocial learning rates (α_other_) and scores on the affective-interpersonal subscale of the Self-Report Psychopathy scale. **(a)** For older adults, levels of psychopathic traits are negatively correlated with prosocial learning rates (*r*_s_=-0.33 [-0.52, -0.11], *p*=0.005, false discovery rate (FDR) corrected *p*=0.03). **(b)** There is no significant relationship for young adults (*r*_s_=0.21 [-0.02, 0.42], *p*=0.07, FDR-corrected *p*=0.22) and the correlation is significantly more negative (Z=3.28, *p*=0.001) for older than young adults. This pattern of results is the same when considering correlations between psychopathic traits and the lack of self-bias in learning (α_other_ - α_self_; not shown). This measure of prosocial learning is also negatively correlated with psychopathic traits in older adults (*r*_s_=- 0.25 [-0.45, -0.02], *p*=0.03) but not younger adults (*r*_s_=0.11 [-0.12, 0.33], *p*=0.36; difference Z=2.15, *p*=0.03). Age-group differences in psychopathic traits and the correlation between α_other_ and psychopathic traits, for older adults only, also remained significant when excluding extreme scores (>3 SDs from the mean) on the psychopathic traits measure (Supplementary Tables 6 & 7). Shaded areas represent 95% confidence intervals.

## Results

We analysed the behaviour of 75 younger adults and 77 older adults who completed the probabilistic reinforcement learning task (Figure 1b), neuropsychological tests, and a measure of psychopathic traits (see Methods). To ensure comparability, older adults with dementia, as diagnosed by the Addenbrooke’s Cognitive Examination (ACE)^49^, were not included in the study. The two age groups were matched on gender (χ^2^(1)=0.45, *p*=0.5) and did not differ in years of education or IQ (Supplementary Table 1). IQ was quantified using age-standardised scores on the Wechsler Test of Adult Reading (WTAR)^50^. We conducted additional analyses controlling for IQ (standardised WTAR score, measured for young and older adults), and memory and attention (memory and attention subscales of the ACE, older adults only). These control analyses showed that our results are not accounted for by general intelligence or executive function (see Methods and Supplementary Information).

### Learning occurs for all recipients for both age groups

We first examined whether participants were able to learn for all three recipients to validate their ability to complete the task. We quantified performance as selecting the option associated with a high chance of receiving reward. Participants in both age groups were able to learn to obtain rewards for themselves, another person, and no one. This was demonstrated through average performance above chance level (50%; all *t*s>15, all *p*s<0.001) and a significant effect of trial number in predicting trial-by-trial performance (all *z*s>4.48, *p*s<0.001) for each separate recipient and age group combination.

### Learning rate depends on who receives reward

Next, to quantify learning, we used computational models of reinforcement learning to estimate learning rates (α) and temperature parameters (β), key indices for the speed by which people update their estimates of reward, and the precision with which they make choices, respectively. Models were fitted using a hierarchical approach and compared using Bayesian model selection as used previously by Huys et al.^51^ and Wittmann et al.^52^ (see Methods). We tested multiple models that varied with respect to whether learning could be explained by shared or separate free parameters across recipient (self, other, no one). Based on our previous results^7^, we examined whether shared or separate learning rates in particular resulted in a better model fit. We used four candidate models:

i. 1α1β: one α for all three recipients & one β for all three recipients;
ii. 3α1β: α_self_, α_other_ & α_no one_, one β;
iii. 2α1β: α_self_ & α_not-self [other + no one]_, one β;
iv. 3α3β: α_self_, α_other_, α_no one_, β_self_, β_other_ & β_no one_ (see Supplementary Table 2).

Initially, we aimed to establish that both our experimental schedule and our models were constructed in a way that allowed us to disentangle recipient-specific learning rates. To this end, we created synthetic choices using simulations based on each of our four models (see Methods). We fitted the models to the data and assured that the best fitting model was the one that had been used to create the data. In such a way, we established model identifiability, both when considering the exceedance probability (Figure 2a and see Methods) and the number of times a model was identified as the best one (Figure 2b). As a second prerequisite for testing for agent-specific learning rates, we performed parameter recovery using our key model of interest, the 3α1β model. Over a wide parameter space, we were able to recover the parameters underlying our choice simulation (Figure 2c).

Having established the models were identifiable and parameters recoverable, we performed Bayesian model selection on the data from our participants. Participant’s choices were best characterised by the 3α1β model. This indicated that the learning process underlying the choices is most accurately captured by assuming separate learning rates for each recipient (α_self_, α_other_ & α_no one_). This model fit the data best (exceedance probability=97%; ΔBIC_int_=122; Figure 2d) and predicted choices well (R^2^=51%; see Methods for further details).

### Higher learning rates are associated with better performance

As a final check of the robustness of our model and to enable clear interpretation of any differences in learning rate between recipients and age groups (c.f. ^53^), we conducted an additional simulation experiment. We simulated data from 10,000 participants using the 3α1β model. This created 30,000 values of α from the three recipient conditions, spanning the full range of possible values from 0 to 1 (see Methods). For each, we quantified the associated performance as the percentage of times the synthetic participant chose the high probability of reward option, averaged across the blocks for the relevant recipient. Plotting the learning rates against performance (Figure 2e) shows that the optimal value of α is approximately 0.55. This is higher than all the values of α found on our task in any recipient condition for either age group. Therefore, higher learning rates were associated with better performance. We further established this link by correlating learning rates and performance in the empirical data from our participants. We found a strong correlation between learning rates and performance overall (*r*_s(150)_ [95% confidence interval]=0.58 [0.46, 0.68], *p*<0.001; Figure 2f) and in each recipient and age group combination (Supplementary Table 3).

### Older adults show a reduced self-bias in learning rates

Next, we used this validated computational model to test our hypotheses as to whether there were group differences in learning rates when learning to reward self, other or no one. Two participants had learning rates for two of the three recipients more than three standard deviations (SDs) above the mean (α_self_ 6.68 & α_no one_ 9.64; α_self_ 7.96 & α_other_ 3.78 SDs above the mean) and were excluded from all analysis of learning rates. We analysed the condition-specific learning rates from our best fitting computational model using a robust linear mixed-effects model (RLMM; see Methods). The RLMM fixed effects were age group (young, older), recipient (self, other, no one), as well as the age group * recipient interaction. While this RLMM includes all three recipient conditions, it generates coefficients (main effect and the interaction with age group) contrasting pairs of recipient conditions – [self vs. other] and [self vs. no one]. These are more interpretable than an omnibus test that would not show which recipient conditions were driving an effect or interaction. We followed up these results with planned comparisons, between the older and younger group in each recipient condition, and between pairs of recipient conditions within each age group.

Across age groups, participants showed a higher learning rate when rewards were for themselves, compared to for another person (recipient [self vs. other]: *b*=-0.024 [- 0.034, -0.014], *z*=-4.79, *p*<0.001). Importantly however, this pattern differed between age groups. The difference between learning rates for self and other was reduced in older compared to younger adults (recipient [self vs. other] * age group interaction: *b*=0.016 [0.002, 0.030], *z*=2.29, *p*=0.02). Between-group comparisons showed older adults learnt more slowly for themselves compared to younger adults (W=3512, Z=- 2.63, r_(150)_=0.22 [0.06, 0.36], *p*=0.009). However, prosocial learning was preserved, with a Bayes factor suggesting strong evidence of no difference in α_other_ between young and older adults (W=3042, Z=-0.86, r_(150)_=0.07 [0.00, 0.24], *p*=0.39, BF_01_=4.26). Within-subject comparisons of α_self_ and α_other_ in each age group showed that young adults had higher learning rates for themselves, relative to another person (V=659, Z=-4.04, r_(75)_=0.47 [0.26, 0.63], *p*<0.001). In contrast, older adults showed no significant difference between learning rates for self and other (V=1150, Z=-1.45, r_(75)_=0.17 [0.01, 0.38], *p*=0.15, BF_01_=1.08).

As expected, across age groups learning was slower for no one than self (recipient [self vs. no one]: *b*=-0.023 [-0.033, -0.013], *z*=-4.57, *p*<0.001). Unlike α_self_ vs. α_other_, learning for self compared to no one did not interact with age group (recipient [self vs. no one] * age group interaction: *b*=0.008 [-0.006, 0.022], *z*=1.15, *p*=0.25). Within-subject comparisons between α_self_ and α_no one_ in each age group showed that both groups learnt preferentially for themselves compared to no one (young adults: V=928, Z=-2.62, r_(75)_=0.30 [0.07, 0.51], *p*=0.009; older adults V=901, Z=-2.76, r_(75)_=0.32 [0.09, 0.53], *p*=0.006). There was no significant difference between the age groups in α_no one_ but also no evidence in support of the null (W=3241, Z=-1.61, r_(150)_=0.13 [0.01, 0.29], *p*=0.11, BF_01_=2.04).

Considering differences in learning between α_other_ and α_no one_, young adults did not differentiate between another person and no one, with strong Bayesian evidence for no difference (V=1533, Z=-0.57, r_(75)_=0.07 [0.00, 0.31], *p*=0.57, BF_01_=5.08). In contrast, older adults had higher learning rates for another person, compared to no one (V=976, Z=-2.37, r_(75)_=0.27 [0.05, 0.49], *p*=0.02). Crucially, this shows that older adults’ lack of differentiation between self and other was not simply because they were insensitive to the recipient condition.

Finally, we also observed an effect of age on both learning rates overall and temperature parameters. Older adults showed slower learning overall compared to younger adults (*b*=-0.019 [-0.028, -0.009], Z=-3.73, *p*<0.001) and higher levels of exploration of choice options (median β young: 0.05, older: 0.19, W=1511, Z=-4.89, r_(150)_=0.40 [0.26, 0.53], *p*<0.001; Supplementary Figure 1).

In summary, older adults prosocial learning was preserved at the same rate as young adults, despite age-related declines in self-relevant learning rates. In other words, young adults showed a self-bias in learning, but older adults distinguished between themselves and others significantly less than the young participants. Only older adults, not young adults, distinguished between rewards for another person and no one.

### Participants perform better for themselves, compared to no one

For completeness, we also tested the effects of recipient and age group on trial-by-trial tendency to pick the high reward stimuli (Figure 3c). In addition to the main effect of trial number (*b*=1.71 [1.29, 2.13], *z*=7.97, *p*<0.001), showing learning, these models revealed older adults chose the high reward option less frequently (mean for young: 80%, older: 64%, *b*=-1.18 [-1.65, -0.70], *z*=-4.84, *p*<0.001), and improved less during the task (trial number * age group interaction *b*=-0.81 [-1.34, -0.27], *z*=-2.95, *p*=0.003) across recipients. When averaging across age groups, performance was better for the self (75%), compared to no one (70%; *b*=-0.36 [-0.05, 0.16], *z*=-2.28, *p*=0.02). However, there was not a significant difference between accuracy for other (72%) and self (*b*=-0.22 [-0.49, 0.05], *z*=-1.63, *p*=0.10), or any significant interactions between age group and recipient (*b*s<0.16, *z*s<0.96, *p*s>0.34).

### Psychopathic traits are lower in older adults and explain variance in prosocial learning

Finally, we examined individual variability in psychopathic traits, considering age-related differences and influence on prosocial learning. Several studies have suggested that individual differences in psychopathic traits can be meaningfully and accurately captured in community samples and often parallel findings in criminal offenders^43^. Critically, psychopathic traits are closely linked to alterations in social behaviour and willingness to help others. Therefore, we also asked participants to complete the Self-Report Psychopathy Scale (SRP-IV-SF)^39^. The SRP is a measure of psychopathic traits in healthy samples that assesses traits linked to clinical psychopathy, such as antisociality and interpersonal affect. The measure robustly captures the latent structure of clinical psychopathy to enable parallels can be drawn between normal individual differences in the community and clinical samples (see Methods). One participant in each age group had missing questionnaire data and are not included in these analyses. Psychopathic traits are consistently divided into two components that this scale measures: core affective-interpersonal traits, which capture lack of empathy and guilt; and lifestyle-antisocial traits, which capture impulsivity and antisocial tendencies. Comparing the two age groups on these scales showed that older participants had significantly lower scores than young participants on both the core affective-interpersonal (young mean=24.36, older mean=21.09, W=3558, Z=-3.15, r_(148)_=0.26 [0.11, 0.40], *p*=0.002) and the lifestyle-antisocial subscales (young mean=22.89, older mean=20.27, W=3471, Z=-2.82, r_(148)_=0.23 [0.09, 0.38], *p*=0.005). These findings suggest that both components of psychopathic traits were reduced in older, compared to younger, adults.

Next, we sought to test our hypothesis that individual differences in core psychopathic traits would explain variability in learning rates, specifically for prosocial learning. We observed a significant negative relationship between α_other_ and core psychopathic traits among older participants (*r*_s(74)_=-0.33 [-0.52, -0.11], *p*=0.005; Figure 4a). Intriguingly, this relationship was significantly more negative (Z=3.28, *p*=0.001) than the equivalent correlation in young adults, which was not significant (*r*_s(74)_=0.21 [-0.02, 0.42], *p*=0.07; Figure 4b). This pattern of results was the same when correlating ‘relative prosocial learning rate’ (the difference between α_other_ - α_self_) with core psychopathic traits. We also conducted control analyses, correlating the same pairs of variables but using partial correlations controlling for β. The negative relationship between prosocial learning (when quantified as α_other_ or α_other_ - α_self_) and psychopathic traits was still present for older adults, showing that the correlations with learning rates were independent of individual choice exploration (all *p*s<0.05; see Supplementary Table 4). The negative relationship between α_other_ and core psychopathic traits for older adults also remained significant after applying false discovery rate (FDR) correction for multiple comparisons across this correlation and the five other age group & recipient combinations (see Supplementary Table 5). Moreover, the finding that no significant correlations were apparent between psychopathic traits and α_self_ or α_no one_ (*p*s>0.15; Supplementary Table 5) suggests a specificity in the relevance of psychopathic traits to prosocial learning in older adults.

Given the age-group differences in levels of psychopathic traits and their correlations with prosocial learning, our final analysis considered whether scores on the core affective-interpersonal psychopathic traits measure mediated the effect of age group on relative prosocial learning rate (α_other_ - α_self_). A standard mediation model (Figure 5a) did not show evidence for a significant mediation. However, as would be predicted if the link between psychopathic traits and prosocial learning depends on age, we found evidence for a moderated mediation. This revealed core psychopathic traits mediated the effect of age group on relative prosocial learning rate for older adults (unstandardised indirect effect=0.006 [0.001, 0.013], *p*=0.008, proportion mediated=0.32) but not for young adults (unstandardised indirect effect=-0.001 [- 0.007, 0.006], *p*=0.65; see Figure 5b for standardised coefficients). In summary, young and older adults differed in levels of psychopathic traits and whether or not psychopathic trait scores explained the extent to which their learning rates were relatively prosocial or self-biased.

**Figure 5.**
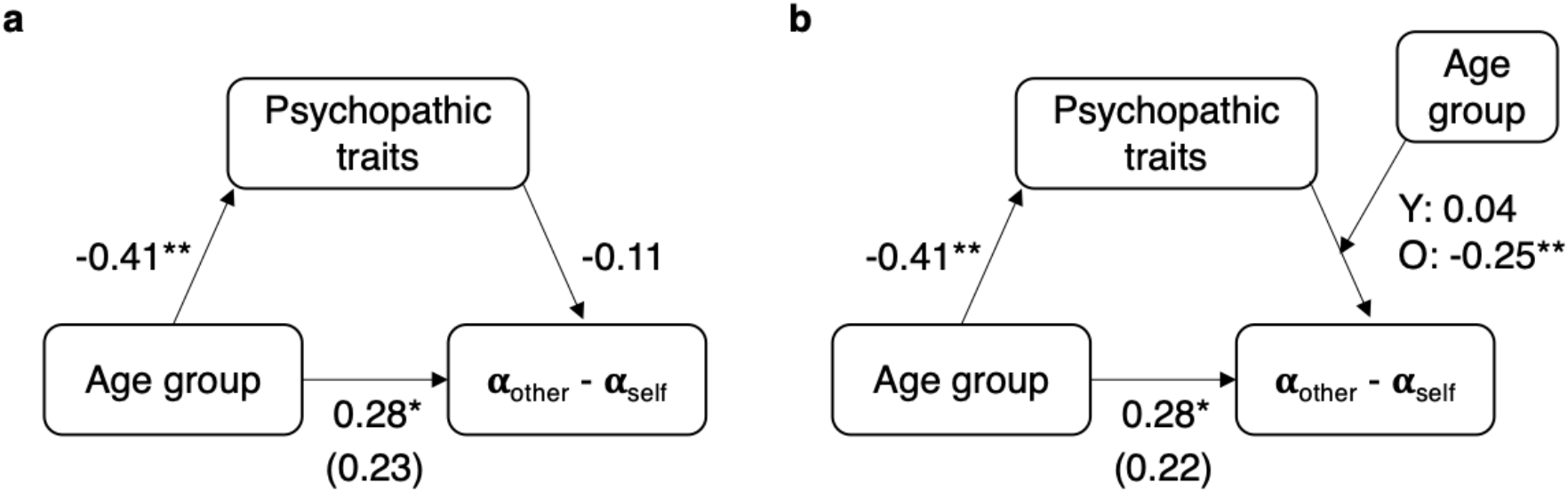
Psychopathic traits mediate the effect of age group on relative prosocial learning (α_other_ - α_self_) for older adults only. **(a)** A standard mediation model does not show evidence that psychopathic traits mediate the effect of age group on relative prosocial learning. Although accounting for psychopathic traits means the significant direct effect of age group on relative prosocial learning (standardised coefficient=0.28, *p*=0.04) becomes non-significant (standardised coefficient=0.23, *p*=0.11), psychopathic traits do not predict relative prosocial learning overall so there is no mediation. **(b)** Evidence of a moderated mediation is revealed when accounting for differences between young and older adults in how psychopathic traits predict relative prosocial learning. Psychopathic traits are a mediator for older but not younger adults. Psychopathic traits are scores on the affective-interpersonal subscale of the Self-Report Psychopathy scale, Y: young, O: older, asterisks represent significant effects (\**p*<0.05, \*\**p*<0.01).

## Discussion

Reinforcement learning is a fundamental process for adaptive behaviour in many species. However, existing studies have largely focused on young people and self-relevant learning, but the decisions we make often occur in a social context^54^ and our actions affect outcomes for others. Here, for the first time, we applied computational models of reinforcement learning to the question of ageing-related changes in self-relevant and prosocial learning. We found a clear decrease in learning rates for self-relevant rewards in older, compared to younger, adults. Intriguingly, despite this reduction in self-relevant learning, learning rates for outcomes that affected others did not differ between older and young adults, with Bayesian analyses supporting no difference. Moreover, not only did older adults have preserved prosocial learning rates, age was also associated with lower psychopathic traits, which were specifically linked to prosocial learning ability in older adults.

Models of learning are a powerful tool for understanding prosocial behaviour. By isolating the learning rate, we can precisely examine the influence of reward history on learning. We robustly replicated previous findings that self-relevant learning can be computationally separated from prosocial learning^7^, with different learning rates for different recipients providing the best explanation of behaviour. Including separate learning rates improved the model fit and, on average across participants, there was a self-bias, learning rates were higher for self-relevant rewards, compared to when someone else received the reward. However, this self-bias was reduced in older adults who showed higher prosocial learning rates, relative to their own self-relevant learning rates, than young adults. As expected, older adults learnt more slowly for themselves than young adults but in the prosocial condition, the learning rates did not significantly differ between the age groups. Bayesian analyses additionally confirmed that prosocial learning was preserved between young and older adults.

As with much research on age-related changes on cognitive and social tasks, our key finding that self-bias in learning is reduced in older adults could be interpreted as due to changes in ability, motivation, or a combination of these factors. Importantly, learning rates were not associated with executive function or an age-standardised measure of intelligence. We also show that our results remain the same after controlling for these measures. These findings suggest that the observed decline in self-relevant learning rates, but preservation of prosocial learning rates, for older adults are not explained by changes in these broad abilities. Considering learning more specifically, a recent comparison of motivation and ability as explanations for ageing-related reductions in model-based strategies during self-relevant learning supported the limited cognitive abilities account^55^. Our finding that learning rates for self-relevant outcomes were reduced in older adults is in line with a degeneration in the neurocognitive systems required for successful learning. Research combining models of learning with neuroimaging and pharmacological manipulations suggests ageing reduces the ability to generate reward prediction errors^18^ (but see^56,57^) due to declines in dopamine functioning^28,58^ (also see^59,60^). Differences in motivation could also be applicable for self-relevant learning as the subjective value of financial outcomes is also likely to decrease in older age, due to changes in wealth across the lifespan^34^.

Our findings suggest that despite declines in learning ability associated with ageing, prosocial learning – learning to help others – is preserved. This finding aligns with an emerging literature showing older adults may be more prosocial and less self-biased than younger adults^29,30,61^. The possibility that relatively preserved prosocial learning is related to increased prosocial motivation is in-line with our observed link between learning rates and psychopathic traits. Psychopathic traits were significantly reduced in our older adult sample, dovetailing with similar previous findings on this trait^47,48^ and broader trait benevolence^33^. Importantly, we found psychopathic traits in older adults negatively correlated with prosocial learning rates. Self-bias in learning rates (i.e. higher learning rate for self compared to other) was most reduced, and even reversed, in the older people lowest on psychopathic traits. Notably, this negative correlation between psychopathic traits and prosocial learning was only found for older adults. This suggests that age-related differences in prosocial learning could be linked to basic shifts in individual traits and motivations over the lifespan, not just to domain-general reductions in cognitive abilities. The idea that social motivations become more influential in learning and decision-making with age has also been suggested based on studies of social rewards such as smiling faces or hypothetical time spent with social partners^62–64^. Taken together, this work suggests that strategies to support healthy ageing might benefit from leveraging potentially preserved social motivations.

Importantly, our task included a control condition where no one benefitted. This was essential to establish that the lack of difference between self and other learning rates for older adults was not simply due to an age-related reduction in the absolute dynamic range as maximum learning rates decrease. Older adults had higher learning rates for both others and themselves, compared to this control condition. In contrast, young adults did not differentiate another person from no one. Older adults therefore showed a relative increase in learning rates that was specific to the prosocial condition. This is also evidence against the idea that lower learning rates in older people are reflective of a general reduced sensitivity to who gets the reward. It is interesting to note that the magnitude of the decrease in self-relevant learning rates associated with being older, compared to younger, is similar to the decrease associated with a young person learning for someone else, compared to themselves. The preservation of prosocial learning rates between age groups may seem at odds with the decreased self-relevant learning rates in our sample of older adults and existing evidence of underlying neurobiological deterioration. However, learning from outcomes for self and other have been linked to distinct regions of the brain in humans, shown though neuroimaging^7,65^, and causally in monkeys with focal lesions^66^.

Taken together, our results add to a growing body of literature suggesting age-related increases in prosocial motivation. If this is the case, the next question is how and why this happens, as there are many possible reasons to be prosocial. For example, prosocial behaviours can be motivated by reputational concerns^67^, the ‘warm glow’ of helping^68–70^, or vicarious reinforcement from positive outcomes for others^71^. In our procedure, we were very careful to prevent reputational concerns influencing learning to help others. Participants underwent an extensive procedure to introduce them to another participant but to hide information about their age and identity. This meant we could assess tendency for prosocial learning in a situation where reputational motivations were excluded, and identity-based influences were controlled. Using a reinforcement learning task, in which performance generates positive outcomes for others also focusses on vicarious rewards from the outcome, rather than warm glow associated with the action of helping. Thus, the increase in prosocial learning rates, relative to self-relevant learning rates, suggests older adults are reinforced by outcomes for others and themselves more similarly than younger adults. Many prosocial measures such as the dictator game^72^ are costly, requiring direct trade-offs between outcomes for oneself and others. This also makes it hard to detect whether changes are in the value of outcomes for oneself, or outcomes for others, or both. In contrast, separating self-relevant learning, prosocial learning, and the control condition allows us to differentiate increases in the value of prosocial outcomes from decreases in the value of outcomes for the self. Our results are consistent with older adults having both decreased self-relevant learning rates (compared to young adults) and increased prosocial learning rates (compared to their performance in the control condition).

Our results also support the advantages of a model-based approach for understanding both prosocial behaviour and ageing. The model-based parameters were more sensitive to the effects of interest than general measures of performance on the task. The model comparison process is able to provide important information about how the learning process takes place, which cannot be revealed from performance measures alone. We showed that learning was best represented by a separate learning rate for each recipient. Moreover, this approach is able to capture additional latent parameters that drive behaviour, such as the inverse temperature, which indexes how closely participants follow the stimulus value. We demonstrate that a single inverse temperature parameter best explained behaviour during learning across recipient conditions, despite learning rates being distinct.

While our procedure and task have many benefits, it is important to also recognise limitations. To test for age-related differences in prosocial learning, we recruited a group of older adults and a group of younger adults. This increases power to detect differences, but we are unable to assess at what age or how quickly changes take place. Also, while our age groups were matched on years of education and IQ, the recruitment from university databases or issues around self-selection may mean that the levels of education and IQ in our sample are not completely representative of the general population. Further studies could include samples recruited entirely from the community and participants across the whole adult lifespan. Future studies should also assess the timescale after which older participants reach similar ceiling levels of performance as younger adults, or whether they are never able to reach the same level of performance. Of similar importance is the question of whether older adults may be able to sustain higher levels of motivation over an extended period of time compared to younger adults, possibly compensating for slower learning speeds. In this study, due to time constraints and the presence of several conditions, we are only able to derive conclusions from a limited time window. Moreover, previous research has suggested that individual differences in empathy – the ability to vicariously experience and understand others’ affect – might relate to differences in prosocial learning^7^. Empathy is positively associated with affective-interpersonal psychopathic traits and might also relate to motivation to help others^73,74^. Further studies could also assess how empathy predicts changes in prosocial learning across the lifespan. Finally, future studies could manipulate the identity of the recipient as we show a preservation in tendency to help others when there are no particular characteristics known about the other person, but these effects might additionally be modulated by factors such as perceived social distance.

To conclude, we find new evidence that despite declines in self-relevant learning in older adults, the ability to learn which actions benefit others is preserved. Moreover, the bias with which people favour self-relevant outcomes is reduced. Not only do older adults have relatively preserved prosocial learning, they also report lower levels of core psychopathic traits that index lack of empathy and guilt, and this trait difference is linked to the changes in prosocial learning. These findings could have important implications for our understanding of reinforcement learning and theoretical accounts of healthy ageing.

## Materials and Methods

### Participants

We recruited 80 young participants and 80 older participants using the same recruitment methods in order to match the samples as closely as possible. Participants were recruited from university databases, which included students and members of the community, social media, and adverts in local newspapers. We excluded anyone who was currently studying or had previously studied psychology and no one took part for course credit. Additional exclusion criteria were previous or current neurological or psychiatric disorder, non-normal or non-corrected to normal vision and, for the older sample, scores on the Addenbrooke’s Cognitive Examination that indicate potential dementia (cut-off score 82)^49^. This sample size gave us 88% power to detect a medium size effect (d=0.5).

Five young and three older participants were excluded due to: diagnosis of a psychiatric disorder at the time of testing (1 young participant); previous study of psychology (2 young participants); and incomplete or low-quality data (2 young and 3 older participants). This left a final sample of 152 participants, 75 young adults (44 females aged 18-36, mean=23.07) and 77 older adults (40 females aged 60-80, mean=69.84). Two older participants were excluded from all analyses involving learning rates due to each having two learning rate estimates more than three standard deviations (SDs) above the mean (for one α_self_ was 6.68 SDs above the mean & α_no_ one 9.64 SDs above the mean; for the second α_self_ was 7.96 & α_other_ 3.78 SDs above the mean). One further participant from each age group was missing data on the SRP measure so are not included in the relevant analyses.

Participants were paid at a rate of £10 per hour plus an additional payment of up to £5 depending on the number of points they earned for themselves during the task. They were also told the number of points that they earned in the prosocial condition would translate into an additional payment of up to £5 for the other participant (see details of the task below). All participants provided written informed consent and the study was approved by the Oxford University Medical Sciences Inter Divisional Research Ethics Committee and National Health Service Ethics.

### Prosocial learning task

The prosocial learning task is a probabilistic reinforcement learning task, with rewards in one of three recipient conditions: for the participant themselves (*self*), for another participant (*other*; prosocial condition), and for *no one* (control condition). Each trial presents two symbols, one associated with a high (75%) probability of gaining points and the other with a low (25%) probability of gaining points. The two symbols were randomly assigned to the left or right side of the screen and selected via a corresponding button press. Participants select a symbol then receive feedback on whether they obtained points or not (see Figure 1b) so learn over time which symbol maximises rewards. The experiment was subdivided into blocks, i.e. 16 trials pairing the same two symbols for the same recipient. Participants completed three blocks, a total of 48 trials, in each recipient condition, resulting in 144 trials overall (see Supplementary Information for trial structure). Blocks for different recipients were pseudo-randomly ordered such that the same recipient block did not occur twice in a row.

On trials in the *self* condition, points translated into increased payment for the participant themselves. These blocks started with “play for you” displayed and had the word “you” at the top of each screen. Blocks in the *no one* condition had “no one” in place of “you” and points were not converted into any extra payment for anyone. In the prosocial *‘other’* condition, participants earned points that translated into additional payment for a second participant, actually a confederate. Participants were told that this payment would be given anonymously, they would never meet the other person, and that the person was not even aware of them completing this task (see Supplementary Information). The name of the confederate, gender-matched to the participant, was displayed on these blocks at the start and on each screen (Figure 1b). Thus, participants were explicitly aware who their decisions affected on each trial. Stimuli were presented using Presentation (Neurobehavioral Systems – https://www.neurobs.com/).

### Questionnaire measures

#### Dementia screening and executive function

Older adults were screened for dementia using the Addenbrooke’s Cognitive Examination (ACE-III)^49^. The ACE examines five cognitive domains; attention, memory, language, fluency and visuospatial abilities. The ACE-III is scored out of 100 and as a screening tool, a cut-off score of 82/100 denotes significant cognitive impairment. We also used scores on the attention and memory domains in control analyses as proxies for executive function in older adults.

#### General intelligence

All participants completed the Wechsler Test of Adult Reading (WTAR)^50^ as a measure of IQ. The WTAR requires participants to pronounce 50 words that have unusual grapheme-to-phoneme translation. This means the test measures reading recognition or existing knowledge of the words, rather than ability to apply rules for pronunciation. The WTAR was developed and standardised with the Wechsler Memory and Adult Intelligence Scales and correlates highly with these measures^75^. Standardisation involves adjusting for healthy age-related differences. The test is suitable for participants aged 16-89, covering our full sample, and scores in older age have been shown to correlate with cognitive ability earlier in life^76^.

#### Psychopathic traits

Participants completed the short form of the Self-Report Psychopathy Scale 4^th^ Edition (SRP-IV-SF)^39^. This scale consists of 29 items, 7 each measuring: interpersonal, affective, lifestyle and antisocial tendencies (plus ‘I have been convicted of a serious crime’). We used the two-factor grouping, summing the core, affective-interpersonal items and separately, the lifestyle-antisocial items for use in analysis. The robust psychometric properties of this measure have been established in community^77^ and offender populations through construct and convergent validity^78^, internal consistency, and reliability^39^.

### Procedure

#### Role assignment

To enhance belief that points earned in the prosocial condition benefitted another person, we conducted a role assignment procedure based on a set-up used in several studies of social decision-making^79,80^. Participants were instructed not to speak and wore a glove to hide their identity. A second experimenter brought the confederate, also wearing a glove, to the other side of the door. Participants only ever saw the gloved hand of the confederate, but they waved to each other to make it clear there was another person there (Figure 1a). The experimenter tossed a coin to determine who picked a ball from the box first and then told the participants which roles they had been assigned to, based on the ball they picked. Our procedure ensured that participants always ended up in the role of the person performing the prosocial learning experiment. Participants were unaware of the age of the other person, but the experimenter used a name for them suggesting their gender was the same as the participant.

#### Task procedure

Participants received instructions for the learning task and how the points they earned would be converted into money for themselves and for the other participant. Instructions included that the two symbols were different in how likely it was that choosing them lead to points but that which side they appeared on the screen was irrelevant. Participants then completed one block of practice trials before the main task and were aware outcomes during the practice did not affect payment for anyone. After the task, participants completed the measure of psychopathic traits and the dementia screening

### Computational modelling

We modelled learning during the task with a reinforcement learning algorithm^16^, creating variations of the models through the number of parameters used to explain the learning rate and temperature parameters in the task^81^. The basis of the reinforcement learning algorithm is the expectation that an action (or stimulus) *a* will provide reward on the following trial. This expected value, *Q*_*t*+1_*(a)* is quantified as a function of current expectations *Q*_*t*_*(a)* and the prediction error *δ*_*t*_, which is scaled by the learning rate α:

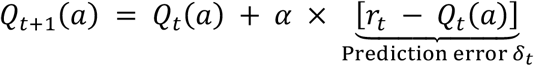

Where *δ*_*t*_, the prediction error, is the difference between the actual reward experienced on the current trial *r*_*t*_ (1 for reward and 0 for no reward) minus the expected reward on the current trial *Q*_*t*_*(a)*.

The learning rate α therefore determines the influence of the prediction error. A low learning rate means new information affects expected value to a lesser extent. The softmax link function quantifies the relationship between the expected value of the action *Q*_*t*_*(a)* and the probability of choosing that action on trial *t* :

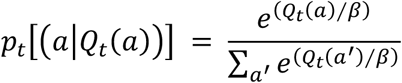

The temperature parameter β represents the noisiness of decisions – whether the participant explores or always chooses the option with the highest expected value. A high value for β means choices seem random as they are equally likely irrespective of the expected value of each option. A low β leads to choosing the option with the greatest expected value on all trials.

### Model fitting

We used MATLAB 2019b (The MathWorks Inc) for all model fitting and comparison. To fit the variations of the learning model (see below) to (real and simulated) participant data we used an iterative maximum a posteriori (MAP) approach previously described^51,52^. This method provides a better estimation than a single-step maximum likelihood estimation (MLE) alone by being less susceptible to the influence of outliers. It does this via implementing two levels: the lower level of the individual subjects and the higher-level reflecting our full sample. For the real participant data, we fit the model across groups to provide the most conservative comparison, so this full sample combined young and older participants.

For the MAP procedure, we initialized group-level Gaussians as uninformative priors with means of 0.1 (plus some added noise) and variance of 100. During the expectation, we estimated the model parameters (α and β) for each participant using an MLE approach calculating the log-likelihood of the subject’s series of choices given the model. We then computed the maximum posterior probability estimate, given the observed choices and given the prior computed from the group-level Gaussian, and recomputed the Gaussian distribution over parameters during the maximisation step. We repeated expectation and maximization steps iteratively until convergence of the posterior likelihood summed over the group, or a maximum of 800 steps. Convergence was defined as a change in posterior likelihood <0.001 from one iteration to the next. Note that bounded free parameters were transformed from the Gaussian space into the native model space via appropriate link functions (e.g. a sigmoid function in the case of the learning rates) to ensure accurate parameter estimation near the bounds. The detailed code for the models and implementation of the fitting algorithm can be found here: https://osf.io/xgw7h/?view_only=bea7b82435b344c5a26f80fd21d8ce19.

### Model comparison

Our hypotheses generated four models to compare that differed in whether the model parameters (α and β) for each participant had one value across recipient conditions or depended on the recipient (self, other and no one). For model comparison, we calculated the Laplace approximation of the log model evidence (more positive values indicating better model fit^82^) and submitted these to a random-effects analysis using the spm_BMS routine^83^ from SPM 8 (http://www.fil.ion.ucl.ac.uk/spm/software/spm8/). This generates the exceedance probability: the posterior probability that each model is the most likely of the model set in the population (higher is better, over .95 indicates strong evidence in favour of a model). For the models of real participant data, we also calculated the integrated BIC (lower is better^51,52^) and R^2^ as additional measures of model fit. To calculate the model R^2^, we extracted the choice probabilities generated for each participant on each trial from the winning model. We then took the squared median choice probability across participants. The 3α1β model had the best evidence on all measures (see Supplementary Table 2).

### Simulation experiments

We used simulation experiments to assess that our experiment allowed us to dissociate models of interest, as well as parameters of interest within the winning model. We simulated data from all four models to establish that our model comparison procedure (see above) could accurately identify the best model across a wide range of parameter values. For this model identifiability analysis, we simulated data from 150 participants, drawing parameters from distributions commonly used in the reinforcement learning literature^84,85^. Learning rates (α) were drawn from a beta distribution (betapdf(parameter,1.1,1.1)) and softmax temperature parameters (β) from a gamma distribution (gampdf(parameter,1.2,5)). We fitted the models to this simulated data set using the same MAP process as applied to the experimental participants’ data and repeated this whole procedure 10 times. By plotting the confusion matrices of average exceedance probability (across the 10 runs; Figure 2a) and how many times each model won (Figure 2b), we show the models are identifiable using our model comparison process.

Our winning model contained 4 free parameters (α_self_, α_other_, α_no one_, β). To assess the reliability of this model and the interpretability of the free parameters, we also performed parameter recovery on simulated data (see Supplementary Information for procedure) as recommended for modelling analyses that use a ‘data first’ approach^81,86^. We simulated choices 1296 times using our experimental schedule and fitted them using MAP. We found strong Pearson’s correlations between the true simulated and fitted parameter values (all *r*s>0.7, see Figure 2c), suggesting our experiment was well suited to estimate the model’s parameters.

Finally, we conducted a principled simulation experiment to identify the optimal learning rate in our task and examine the link between learning rates and performance. We simulated data from 10,000 participants with learning rates (α) and softmax temperature parameters (β) drawn from beta and gamma distributions respectively, as described above. We bounded the β parameters at 0 and 0.3 to reflect the range shown by our participants. In line with our winning model, simulated participants had a separate learning rate for each recipient condition, and these spanned the full range of possible α values from 0 to 1. This generated 30,000 learning rate values (10,000 participants and three conditions). For each we quantified performance as the proportion of times the simulated participant chose the ‘correct’ high reward option in the relevant condition. Results showed that average performance improved as learning rates increased, up to learning rates of approximately 0.55 (Figure 2e). Crucially, this optimal alpha was above the highest learning rate shown by any of our participants in any recipient condition, meaning a higher learning rate on our task was associated with better performance.

### Statistical analysis

Analysis of group and recipient differences in the fitted model parameters and behavioural data was run in R^87^ with R Studio^88^. We used a robust linear mixed-effect model (RLMM; rlmer function; robustlmm package^89^ to predict learning rates and generalised linear mixed-effects models (glmer function; lme4 package^90^) for the trial-by-trial data (binary outcome of choosing the high vs. low reward option). We used (robust / generalised) linear mixed-effects models as these account for the within-subject nature of the recipient manipulation and do not rely on parametric assumptions. Additionally, unlike an anova model or omnibus test, an RLMM generates coefficients (with confidence intervals and significance values) for terms that compare pairs of factor levels (for example self vs. other). This approach is more informative than an anova when the factor has more than two levels, which is our case as the recipient factor has three levels (self, other, no one). Each linear mixed-effects model had fixed effects of age group, recipient (self, other, no one), and their interaction, plus a random subject-level intercept. Analysis of trial-by-trial choices also included trial number in the fixed terms, interacting with recipient and group (including the three-way interaction), and in the random terms, interacting with recipient. In the analysis of learning rates controlling for IQ, standardised scores on the WTAR were also included as a fixed term (Supplementary Table 8). Correlations of learning rates with psychopathic traits (Supplementary Table 4 & 5) and neuropsychological measures (Supplementary Table 9) were calculated with Spearman’s Rho nonparametric tests. To control for IQ and executive function in the associations between older adults’ prosocial learning rates and psychopathic traits, we ran partial correlations each controlling for one of WTAR, ACE memory and ACE attention scores (Supplementary Table 10).

For simple and post hoc comparisons, we used two-sided nonparametric tests as outcome variables violated normality assumptions. Effect sizes and confidence intervals for paired and independent nonparametric comparisons were calculated using the *cohens_d* and *wilcox_effsize* functions respectively from the rstatix package^91^. Bayes factors (BF_01_) for non-significant results were calculated using nonparametric paired and independent t-tests in JASP^92^ with the default prior. BF_01_ corresponds to how many times more likely the data are under the null hypothesis of no difference than under the alternative hypothesis that there is a difference. A BF_01_ larger than 3 (equal to BF_10_ less than 1/3) is considered substantial evidence in favour of the null hypothesis whereas a BF_01_ between 1/3 and 3 indicates the data cannot clearly differentiate between hypotheses^93^. Median learning rates and their standard errors for plotting were calculated using bootstrapping with 1,000 samples.

For the mediation analysis, we used the *mediate* function (mediation package^94^) combined with robust linear models (*rlm* function, MASS package^95^). This method estimates the unstandardised indirect effects, with 95% confidence intervals and significance, through a bootstrapping procedure with 10,000 bootstrapped samples. The outcome in the mediation model was relative prosocial learning rate (α_other_ - α_self_), the predictor was age group, and the mediator was psychopathic traits on the core affective-interpersonal subscale of the SRP-IV-SF. We calculated two mediation models. The first, a standard mediation model, included an indirect path from age group to relative prosocial learning rate via psychopathic traits as a mediator (Figure 5a). The second included an interaction between age group and psychopathic traits in predicting relative prosocial learning rate to allow the possibility of a moderated mediation^96^. Specifically, this type of moderated mediation examines whether the effect of the mediator (in this case psychopathic traits) on the outcome (α_other_ - α_self_) is moderated by the predictor (age group)^97,98^. In other words, psychopathic traits could be a mediator for one age group but not the other (Figure 5b).

## Data availability

Data are available at: https://osf.io/xgw7h/?view_only=bea7b82435b344c5a26f80fd21d8ce19.

## Code availability

Code for modelling and analysis is available at: https://osf.io/xgw7h/?view_only=bea7b82435b344c5a26f80fd21d8ce19.

## Acknowledgements

This work was supported by a Medical Research Council Fellowship (MR/P014097/1), a Christ Church Junior Research Fellowship, and a Christ Church Research Centre Grant to PL; a Wellcome Trust Principal Fellowship to MH; NIHR Biomedical Research Centre, Oxford. The Wellcome Centre for Integrative Neuroimaging is supported by core funding from the Wellcome Trust (203139/Z/16/Z).

We are grateful to Craig Neumann for assistance with the Self-Report Psychopathy Scale. We are also grateful to the many people who acted as confederates for us during the study.

## Author contributions

PL designed the study. AA, LH, DD & PL collected the data. JC, MW & PL analysed the data. JC, MW, MH & PL wrote the paper.

