## Supplementary Information for "Ageing disrupts reinforcement learning whilst learning to help others is preserved"

### Experimental procedures

#### *Participant instructions*

"In this experiment you will see a pair of symbols on each trial and you need to select one of them.

You will receive points for some of your choices, which will be converted into money at the end of the experiment, so the more points you get the more extra money you will earn. The two symbols are not the same in terms of how often they give you points: some of the symbols will give you points more often than others. Each symbol has its own meaning, regardless of where it appears on the screen (so left and right are not important) or when it occurs in the task.

You will play the task in three recipients, for yourself, for the other participant, and for no one. When you are playing for yourself, you will receive any money you win. When you are playing for the other participant, they will receive any money you win for them. However, the other participant won't know how much money you earn from them until they leave the experiment (you will leave at different times, and the money you earn for the other participants will be given to them in a sealed envelope), and they do not know that you are performing a task where you could win extra money for them. When you play for no one, neither of you will receive any of the money.

Respond using the left and right arrow keys to make your selection. Try to respond as quickly as possible, you have about 2 seconds to make your choice."

#### *Trial structure*

At the start of each block, the recipient was displayed for 2,000ms in the form "Play for [you / confederate name / no one]" (Figure 1b). Then two abstract symbols, in fact letters from the Agathodaimon font, were presented either side of a fixation cross. Participants were required to select one within 3,000ms by pressing the left or the right key. If they did not respond, the word "missed" in red text appeared. If the participant did select an outcome, it was displayed for a further 300ms before a delay of 2,500ms showing a fixation cross. The outcome of their choice, either 100 points or 0 points, was then revealed (800ms) before another fixation (1000ms) period and the next trial, showing the same two symbols again. On all screens other than during fixations, the recipient was displayed above the symbols or the outcome.

A new pair of symbols was used on each block of 16 trials. There were three blocks for each recipient meaning 48 self trials, 48 other trials, and 48 no one trials, so 144 trials in total. Blocks were presented in one of 6 possible orders that ensured the same recipient condition never occurred twice in a row. Participants were randomly assigned to an order. Which side each symbol was presented on was counterbalanced to prevent action learning. The instructions to participants also stated that the location of the symbol was meaningless.

**Parameter recovery**

We simulated choices for our winning model using the trial schedule used in the task (participants were randomly allocated to one of six recipient block orders but as each block is fitted separately this is not relevant for the parameter recovery). We combined a range of parameter values (1296 combinations in total) from a grid of values (0, 0.2, 0.4, 0.6, 0.8, 1). We added noise to each parameter for each simulated participant (using a standard normal distribution multiplied by 0.05) to improve our coverage of possible parameter values. All parameters were bounded at 0 and 1 as learning rates are always in this range and the range of temperature parameters in our sample was similar to that of the learning rates. We then refit the simulated choices using the same MAP process as applied to the experimental participants' data.

In some situations, it is important to also run parameter recovery on the other models to show that the behavioural effect of interest cannot be retrieved[1]. However, we only applied parameter recovery to the winning model as by definition the models with 1α or 2α cannot show the differences in learning rates we find between all three recipient conditions. Differences in β values between recipients were not of interest for our hypotheses.

**Comparing model fit between groups**

Overall, the results of our modelling suggest that a model including different learning rates for each recipient condition explains the data better than a model with a single learning rate across self, prosocial, and no one recipients. We then compared the model fit between age groups. A Wilcoxon between-groups comparison of $R^2$ showed the fit was significantly better for the young than the older participants (median $R^2$ for young: 72%, older: 31%,W=4218, Z=-4.90, $r_{(150)}$=0.40 [0.25, 0.53], $p$<0.001).

To further investigate this difference in model fit, we used the mbb-vb-toolbox[2] (http://mbb-team.github.io/VBA-toolbox/) for random-effects Bayesian model comparisons based on the subject-level log model evidence for each age group separately. This allowed us to estimate the model frequency or probability that the model generated a given subject's data. Here we excluded the two older participants who were removed from the analysis of learning rates due to being outliers.

This output and the exceedance probabilities revealed that the 3α1β model was best for young participants (exceedance probability=1, expected frequency=0.78). However, for older adults, the 1α1β was the winning model (exceedance probability =0.98, expected frequency=0.58) and the expected frequency of the 3α1β model in older adults was 0.36. This result further supports our conclusions in the main manuscript. It indicates that the increase in self learning rate, compared to other learning rate, is abolished with older age. Using parameters from a hierarchical fit of the same, 3α1β model across all participants provides the most conservative test of age-related differences in prosocial learning rates.

### Supplementary results

**Supplementary Table 1.** *Comparison of education and intelligence scores between age groups*

| | | Years of education | IQ |
|---|---|---|---|
| **Young** | Mean (SD) | 14.76 (1.71) | 118 (7.55) |
| | Range | 13-17 | 99-129 |
| **Older** | Mean (SD) | 15.07 (2.69) | 119 (7.57) |
| | Range | 6-20 | 80-128 |
| **Difference** | W | 2511 | 2700 |
| | Z | -1.19 | -0.69 |
| | $r_{(152)}$ | 0.10 | 0.06 |
| | [95% CI] | [0.01, 0.27] | [0.00, 0.23] |
| | *p* | 0.24 | 0.49 |

Note. IQ: standardised score from the Wechsler Test of Adult Reading, SD: standard deviation, W: between-group Wilcoxon t-test statistic, Z: standardised measure of the difference, $r_{(152)}$: effect size$_{(n)}$ of the difference, 95% CI: 95% confidence interval around the effect size, *p*: *p*-value from the Wilcoxon t-test.

**Supplementary Table 2.** *Model comparison results*

| | | $\alpha$ | | | $\beta$ | | | lme | BICint | XP | $R^2$ |
|---|---|---|---|---|---|---|---|---|---|---|---|
| | | s | o | n | s | o | n | | | | |
| Model 1 | $1\alpha\ 1\beta$ | | $\alpha$ | | | $\beta$ | | -10950 | 21889 | 0.03 | 51% |
| Model 2 | $3\alpha\ 1\beta$ | $\alpha_s$ | $\alpha_o$ | $\alpha_n$ | | $\beta$ | | -10801 | 21607 | 0.97 | 51% |
| Model 3 | $2\alpha\ 1\beta$ | $\alpha_s$ | $\alpha_{not\text{-}self}$ | | | $\beta$ | | -10857 | 21729 | 0.00 | 49% |
| Model 4 | $3\alpha\ 3\beta$ | $\alpha_s$ | $\alpha_o$ | $\alpha_n$ | $\beta_s$ | $\beta_o$ | $\beta_n$ | -10977 | 21947 | 0.00 | 50% |

Note. s: self, o: other, n: no one, lme: log model evidence, BICint: integrated Bayesian Information Criterion, XP: exceedance probability.

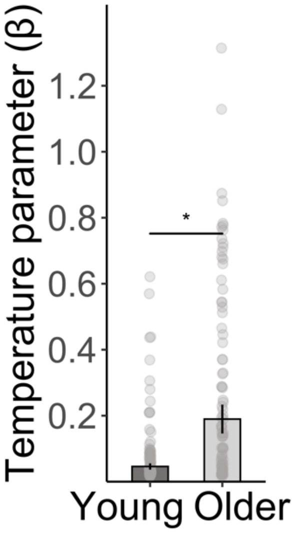


**Supplementary Figure 1. Comparison of inverse temperature parameters (β) between age groups**. Older adults had significantly higher β parameters than young adults, suggesting age is associated with less consistency in choices during the task. Bars show group median, error bars are standard error of the median, the asterisk represents a significant difference from a between-group Wilcoxon t-test ($p$<0.001); n=152 (75 young, 77 older).

**Supplementary Table 3.** *Positive correlations between learning rates and performance*

| | Young | | | Older | | |
|---|---|---|---|---|---|---|
| | $\alpha_{self}$ | $\alpha_{other}$ | $\alpha_{no\ one}$ | $\alpha_{self}$ | $\alpha_{other}$ | $\alpha_{no\ one}$ |
| $r_s$ | 0.46 | 0.66 | 0.65 | 0.68 | 0.66 | 0.61 |
| $p$ | <0.001*** | <0.001*** | <0.001*** | <0.001*** | <0.001*** | <0.001*** |

Note. Performance quantified as the percentage of trials on which participants choose the high reward option. Asterisks represent significance (***$p$<0.001).

**Supplementary Table 4.** *Correlations between prosocial learning and affective-interpersonal psychopathic traits for older adults, controlling for β*

| | $\alpha_{other}$ | | $\alpha_{other}$ - $\alpha_{self}$ | |
|---|---|---|---|---|
| | Standard | Partial | Standard | Partial |
| $r_s$ | -0.33 | -0.35 | -0.25 | -0.25 |
| $p$ | 0.005* | 0.002* | 0.03* | 0.03* |

Note. Correlations between prosocial learning rate and affective interpersonal SRP score, partial correlations controlling for the inverse temperature parameter (β). Asterisks represent significance (*$p$<0.05).

**Supplementary Table 5.** *Correlations between learning rates and affective-interpersonal psychopathic traits*

| | Young | | | Older | | |
|---|---|---|---|---|---|---|
| | $\alpha_{self}$ | $\alpha_{other}$ | $\alpha_{no\ one}$ | $\alpha_{self}$ | $\alpha_{other}$ | $\alpha_{no\ one}$ |
| $r_s$ | -0.02 | 0.21 | -0.13 | 0.17 | -0.33 | -0.01 |
| *p* | 0.85 | 0.07 | 0.27 | 0.16 | 0.005* | 0.94 |
| FDR *p* | 0.94 | 0.22 | 0.40 | 0.31 | 0.03* | 0.94 |

Note. FDR: false discovery rate correction. Asterisks represent significance (*$p$<0.05).

**Supplementary Table 6.** *Comparisons of psychopathic traits between groups excluding extreme scores*

| Subscale | Young mean | Older mean | W | Z | $r_{(df)}$ [95% CI] | *p* |
|---|---|---|---|---|---|---|
| Affective-interpersonal | 23.65 | 20.19 | 3411 | -3.28 | $r_{(142)}$=0.27 [0.12, 0.42] | 0.001** |
| Lifestyle-antisocial | 22.55 | 19.65 | 3395 | -3.04 | $r_{(143)}$=0.25 [0.10, 0.40] | 0.002** |

Note. Extreme scores on the psychopathic traits measure (Self-Report Psychopathy Scale) were defined as those more than 3 standard deviations from the mean, the exclusion criteria applied to learning rates for all analyses. Comparisons are between-group Wilcoxon t-tests. Asterisks represent significance (**$p$<0.01).

**Supplementary Table 7.** *Correlations between learning rates and affective-interpersonal psychopathic traits excluding extreme scores*

| | Young | | | Older | | |
|---|---|---|---|---|---|---|
| | $\alpha_{self}$ | $\alpha_{other}$ | $\alpha_{no\ one}$ | $\alpha_{self}$ | $\alpha_{other}$ | $\alpha_{no\ one}$ |
| $r_s$ | -0.03 | 0.15 | -0.07 | 0.12 | -0.31 | 0.00 |
| *p* | 0.79 | 0.20 | 0.58 | 0.30 | 0.008* | 0.97 |
| FDR *p* | 0.95 | 0.60 | 0.87 | 0.60 | 0.05* | 0.97 |

Note. Extreme scores on the psychopathic traits measure (Self-Report Psychopathy Scale affective-interpersonal subscale) were defined as those more than 3 standard deviations from the mean, the exclusion criteria applied to learning rates for all analyses. FDR: false discovery rate correction. Asterisks represent significance (*$p$<0.05).

**Supplementary Table 8.** *Fixed-effect parameter z-values from robust mixed-effects models predicting learning rates, main model and controlling for general intelligence*

| | Main model | Controlling for IQ |
|---|---|---|
| Intercept | 25.45 *** | 4.35 *** |
| IQ | | -0.43 |
| Recipient [self vs. other] | -4.79 *** | -4.80 *** |
| Recipient [self vs. no one] | -4.57 *** | -4.57 *** |
| Age group [young vs. older] | -3.73 *** | -3.72 *** |
| Recipient [self vs. other] * age group | 2.29 * | 2.29 * |
| Recipient [self vs. no one] * age group | 1.15 | 1.15 |

Note. IQ: standardised score from the Wechsler Test of Adult Reading. Asterisks represent significance (**p*<0.05, ***p*<0.01, ****p*<0.001).

**Supplementary Table 9.** *Correlations between learning rates and neuropsychological measures*

| Measure | | Young | | | Older | | |
|---|---|---|---|---|---|---|---|
| | | $\alpha_{self}$ | $\alpha_{other}$ | $\alpha_{no\ one}$ | $\alpha_{self}$ | $\alpha_{other}$ | $\alpha_{no\ one}$ |
| **IQ** | $r_s$ all participants | 0.04 | | | | | |
| | $r_s$ each group | | 0.03 | | | 0.04 | |
| | $r_s$ each agent | 0.13 | -0.20 | -0.09 | -0.08 | 0.08 | 0.15 |
| **Memory** | $r_s$ each group | | | | | 0.10 | |
| | $r_s$ each agent | | | | -0.03 | -0.00 | 0.12 |
| **Attention** | $r_s$ each group | | | | | 0.15 | |
| | $r_s$ each agent | | | | 0.09 | -0.04 | 0.20 |

Note. IQ: standardised score from the Wechsler Test of Adult Reading, Memory: Addenbrooke's Cognitive Examination (ACE) domain score, Attention: ACE domain score. ACE only measured in older adults. No correlations significant (*p*s>0.05).

**Supplementary Table 10.** *Correlations between prosocial learning and affective-interpersonal psychopathic traits for older adults, controlling for general intelligence and executive function*

| | $\alpha_{other}$ | | | | $\alpha_{other}$ - $\alpha_{self}$ | | | |
|---|---|---|---|---|---|---|---|---|
| | Stnd | Partial | | | Stnd | Partial | | |
| | | IQ | Memory | Attention | | IQ | Memory | Attention |
| $r_s$ | -0.35 | -0.32 | -0.33 | -0.33 | -0.25 | -0.24 | -0.25 | -0.25 |
| *p* | 0.002 | 0.006 | 0.004 | 0.004 | 0.03 | 0.04 | 0.04 | 0.03 |

Note. Stnd: standard correlations between prosocial learning rate and affective interpersonal Self-Report Psychopathy score. Partial correlations controlling for individual scores on IQ: standardised score from the Wechsler Test of Adult Reading, Memory: Addenbrooke's Cognitive Examination (ACE) domain score, Attention: ACE domain score. All correlations significant ($p$<0.05).
